# MGSE regulates crosstalk from the mucin pathway to the TFE3 pathway of the Golgi stress response

**DOI:** 10.1101/710616

**Authors:** Mohamad Ikhwan Jamaludin, Sadao Wakabayashi, Kanae Sasaki, Ryota Komori, Hirotada Kawamura, Hayataka Takase, Miyu Sakamoto, Hiderou Yoshida

## Abstract

The Golgi apparatus is an organelle where membrane or secretory proteins receive post-translational modifications such as glycosylation and sulfation, after which the proteins are selectively transported to their final destinations through vesicular transport. When the synthesis of secretory or membrane proteins is increased and overwhelms the capacity of the Golgi (Golgi stress), eukaryotic cells activate a homeostatic mechanism called the Golgi stress response to augment the capacity of the Golgi. Four response pathways of the Golgi stress response have been identified, namely the TFE3, CREB3, HSP47, and proteoglycan pathways, which regulate the general function of the Golgi, apoptosis, cell survival, and proteoglycan glycosylation, respectively. Here, we identified a novel response pathway that augments the expression of glycosylation enzymes for mucins in response to insufficiency in mucin-type glycosylation in the Golgi (mucin-type Golgi stress), and we found that expression of glycosylation enzymes for mucins such as GALNT5, GALNT8, and GALNT18 was increased upon mucin-type-Golgi stress. We named this pathway the mucin pathway. Unexpectedly, mucin-type Golgi stress induced the expression and activation of TFE3, a key transcription factor regulating the TFE3 pathway, suggesting that the activated mucin pathway sends a crosstalk signal to the TFE3 pathway. We identified an enhancer element regulating transcriptional induction of TFE3 upon mucin-type Golgi stress, and named it the mucin-type Golgi stress response element, of which consensus was ACTTCC(N9)TCCCCA. These results suggested that crosstalk from the mucin pathway to the TFE3 pathway has an important role in the regulation of the mammalian Golgi stress response.

## Introduction

Each organelle in eukaryotic cells has a certain capacity to execute its function. When this capacity becomes insufficient, homeostatic mechanisms will automatically augment the capacity in accordance with cellular needs, which is indispensable for maintaining cellular function and is termed organelle autoregulation (Sasaki and Yoshida, 2015). In principle, organelle autoregulation consists of four factors: (1) sensors that detect insufficiency of organelle function, (2) transcription factors that upregulate the expression of genes related to organelle function, (3) enhancer elements to which the transcription factors bind, and (4) target genes whose expression is augmented by organelle autoregulation, resulting in upregulation of organelle capacity.

Organelle autoregulation of the endoplasmic reticulum (ER) has been analyzed extensively and is known as the ER stress response (Karagoz *et al.*, 2019; Kimata and Kohno, 2011; Mori, 2015; Volmer and Ron, 2015; Yoshida, 2007). In the ER, secretory and membrane proteins are synthesized and folded with the assistance of ER chaperones (Gething, 1997). Proteins that cannot be folded are degraded by a mechanism called ER-associated degradation (ERAD) (Wu and Rapoport, 2018). When the synthesis of secretory and membrane proteins increases and overwhelms the folding capacity of the ER, unfolded and misfolded proteins accumulate in the ER (ER stress), leading to ER stress-induced apoptosis. To cope with ER stress, mammalian cells activate three response pathways of the ER stress response: the PERK, ATF6, and IRE1 pathways. Activation of sensors of these pathways (PERK, pATF6(P), and IRE1) transduces a signal to transcription factors such as ATF4, pATF6α(N), pATF6β(N), and pXBP1(S), and induces the expression of ER-related genes encoding ER chaperones and ERAD components, leading to the augmentation of ER capacity.

Recently, organelle autoregulation of the Golgi apparatus (the Golgi stress response) has been elucidated (Sasaki and Yoshida, 2019). To date, at least four response pathways of the Golgi stress response have been revealed in mammalian system, namely the TFE3, CREB3, HSP47, and proteoglycan pathways (Taniguchi and Yoshida, 2017). TFE3 is a key transcription factor regulating the TFE3 pathway. In normal growth conditions, TFE3 is phosphorylated and retained in the cytoplasm. Upon Golgi stress, TFE3 is dephosphorylated and translocated into the nucleus and activates the transcription of target genes by binding to the Golgi apparatus stress response element (GASE) enhancer element (Taniguchi *et al.*, 2015; Taniguchi *et al.*, 2016). The target genes of the TFE3 pathway include N-glycosylation enzymes, Golgi structural proteins, and vesicular transport component (Oku *et al.*, 2011), suggesting that the TFE3 pathway augments general function of the Golgi. The sensor molecule for TFE3 pathway remains unclear. The CREB3 pathway was identified by Reiling and colleagues (Reiling *et al.*, 2013), in which a key transcription factor is CREB3. CREB3 is a transmembrane protein located in the ER membrane and functions as a sensor for Golgi stress, whereas upon Golgi stress CREB3 is translocated from the ER to the Golgi and is cleaved by S1P and S2P proteases. The cytosolic portion of cleaved CREB3 translocates to the nucleus and activates transcription of ARF4, resulting in apoptosis. In contrast, the HSP47 pathway increases the expression of HSP47, an ER chaperone specialized for collagen folding and maturation, and suppresses Golgi stress-induced apoptosis (Miyata *et al.*, 2013). However, the sensor, transcription factor, and the enhancer element of the HSP47 pathway remain to be identified. We reported recently on the proteoglycan pathway, which increases the expression of glycosylation enzymes for proteoglycans when glycosylation of proteoglycans becomes insufficient (proteoglycan-type Golgi stress) (Sasaki *et al.*, 2019). PGSE-A and PGSE-B are enhancer elements regulating the proteoglycan pathway, although the transcription factors and sensors are unidentified. The proteoglycan pathway seems to be a cell-type-specific response pathway and to be upregulated in cells producing proteoglycans including chondrocytes and reactive astrocytes (Sakamoto and Kadomatsu, 2017).

BenzylGalNAc (BG) is a Golgi stress inducer that has been utilized for activation of the HSP47 pathway (Miyata et al., 2013). BG is an inhibitor of mucin-type glycosylation in the Golgi (Huet *et al.*, 1998; Kuan *et al.*, 1989), and it causes the accumulation of mucin core proteins in the Golgi (Huet et al., 1998). Thus, we speculated that if mucin-producing cells are treated with BG, the capacity of mucin glycosylation is suppressed, resulting in accumulation of less glycosylated mucin core proteins in the Golgi (mucin-type Golgi stress), and that cells may activate an unknown response pathway of the Golgi stress response to upregulate the expression of glycosylation enzymes for mucins, namely the mucin pathway. Here, we investigated whether mammalian cells have the mucin pathway of the Golgi stress response.

## Materials and Methods

### Cell culture and transfection

HT29 (human colon adenocarcinoma cell) and HeLa cells were cultured in Dulbecco’s modified Eagle’s medium (DMEM) (Wako, Osaka, Japan) supplemented with 10% fetal bovine serum (FBS) and kept in a humidified atmosphere of 5% CO_2_ at 37°C (Yoshida *et al.*, 2009). The calcium phosphate or PEI max methods were employed to transfect plasmid DNAs into cells, treated with 10 mM BG for 48 h, washed with PBS and subjected to luciferase assays, immunoblotting, or immunofluorescence (Yoshida *et al.*, 2006).

### Construction of plasmids

Corresponding regions of the human TFE3 promoter were amplified from human genomic DNA and cloned into the *Bgl*II site upstream of the firefly luciferase gene in a pGL4 basic vector (Promega, Fitchburg WI). A 4×GASE plasmid was constructed in the previous report (Oku et al., 2011). Point mutants were made by site-directed mutagenesis using a QuikChange Site-Directed Mutagenesis kit (Stratagene, CA) (Yoshida *et al.*, 2001).

### Quantitative RT-PCR and luciferase assay

Quant-Studio 6 Flex Real-Time PCR System (Thermo Fisher Scientific, Waltham, MA) and PrimeScript RT reagent kit with gDNA Eraser and SYBR Premix Ex Taq II (T1i RNase H Plus) (TaKaRa, Otsu, Japan) were utilized to carry out quantitative RT-PCR (qRT-PCR) (Uemura *et al.*, 2013). Nucleotide sequences of primer pairs used here were as follows: TFE3 (ACTGGGCACTCTCATCCCTAAGTC and TTCAGGATGGTGCCCTTGTTC), HSP47 (AAGAGCAGCTGAAGATCTGGATG and GTCGGCCTTGTTCTTGTCAATG), GALNT5, (GGCCTGTCCAGTAATCGAAGTCA and AAAGTTCATGGGCCACACAAAGA), GALNT8 (AACCATGCTCCAAGGCAGCTA and CATCTCCAGACACCGCTTGGTA) and GALNT18 (TGCCTGACCTCAGACCCCAGT and TGTCATCCACCAGAATGATCTCC). Dual-luciferase reporter assays were performed according to previous reports and Renilla luciferase was utilized as standard for transfection efficiency measurement (Komori *et al.*, 2012).

### Immunoblotting and Immunocytochemistry

Immunoblotting and immunocytochemistry were performed as described previously (Yoshida et al., 2009). Anti-Giantin and anti-GM130 antisera were purchased from COVANCE (Princeton, NJ) (#PRB-114C) and Becton Dickinson and Company (Franklin Lakes, New Jersey) (#610823), respectively. Anti-TFE3 antiserum was prepared in the previous study (Taniguchi et al., 2015).

### Microarray analyses and RNA sequencing with a next-generation DNA sequencer

Total RNA was prepared from HT29 cells treated with 10 mM BG for 48 h utilizing RNeasy Mini kit (Qiagen, Germany), and was subjected to microarray and RNA sequencing analyses as described previously (Sasaki et al., 2019).

## Results

### Expression of genes encoding enzymes for mucin-type glycosylation is elevated upon BG treatment

BG is an inhibitor of C1GALT1 (a key enzyme for mucin-type glycosylation) (Kuan et al., 1989), which can make mucin-type glycosylation insufficient (mucin-type Golgi stress), and has been used to activate the HSP47 pathway of the Golgi stress response (Miyata et al., 2013). Upon treatment with BG, it was reported mucin-type O-glycosylation in the Golgi is severely inhibited, and that the level of HSP47 mRNA was increased in Colo 205 and NIH3T3 cells in order to suppress Golgi stress-induced cell death. Prompted by a previous study, we analyzed the expression of HSP47 mRNA in HT29 cells using qRT-PCR to confirm that BG can induce mucin-type Golgi stress in HT29 cells. As shown in Fig. 1A, we found that the expression of HSP47 was elevated after BG treatment for 48 h (lanes 1 and 2). This finding indicated that BG induced Golgi stress and stimulated an increase in HSP47 mRNA in our experimental system.

**Fig. 1.**
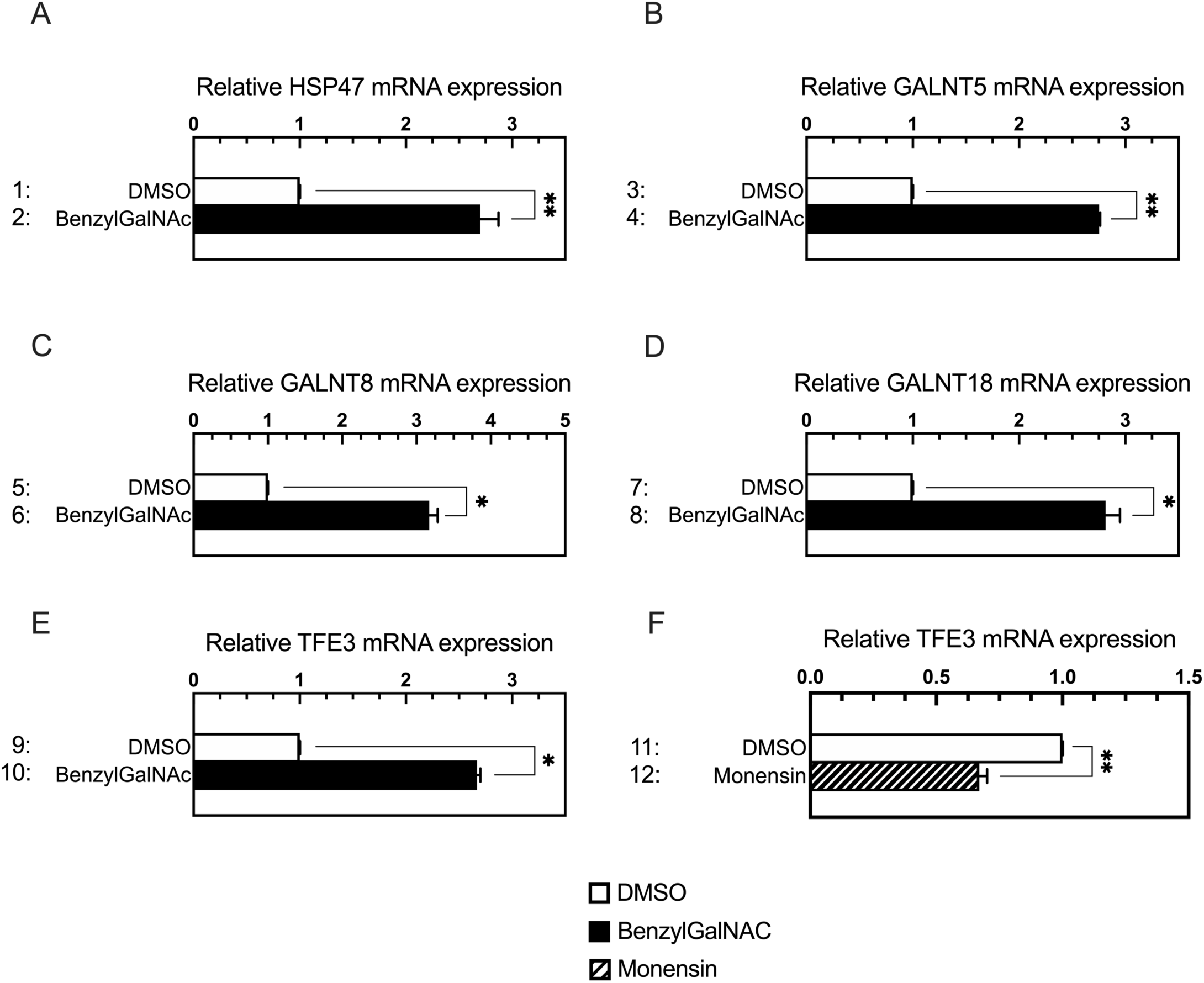
Effect of BG on the expression of HSP47, GALNT8, GALNT5, GALNT18, and TFE3 mRNAs. Total RNA prepared from HT29 cells treated with 10 mM BG for 48 h or 0.3 μM monensin were subjected to qRT-PCR experiments (A, B, D, E, and F) or to RNA sequencing (C) to evaluate levels of the indicated mRNAs. HT29 cells treated with DMSO was used as the control. Values are means ± SE of two and three independent experiments. ***, P < 0.001; **, P < 0.01; *, P < 0.05.

We assumed that BG treatment may induce the expression of enzymes for mucin-type glycosylation as well as HSP47, because BG causes insufficiency of mucin glycosylation (mucin-type Golgi stress), and we examined whether genes encoding mucin-type glycosylation enzymes are induced upon BG treatment. When HT29 cells were treated with BG for 48 h and subjected to RNA sequencing and qRT-PCR analyses, the expression levels of GALNT5, GALNT8, and GALNT18 were induced (Fig. 1B-D). These results strongly suggested that the expression of enzymes for mucin-type glycosylation is augmented by mucin-type Golgi stress (insufficiency of mucin glycosylation). We named this novel pathway the mucin pathway.

### Transcription of TFE3 mRNA is increased by BG treatment

TFE3 is a transcription factor regulating the TFE3 pathway of the mammalian Golgi stress response (Taniguchi et al., 2015). It has been revealed that TFE3 is activated by dephosphorylation at Ser108, although the expression of TFE3 mRNA was not changed upon treatment with activators of the TFE3 pathway, such as monensin and nigericin, and expression of a dominant negative form of GCP60. Surprisingly, we found that the expression of TFE3 mRNA was increased by BG treatment (Fig. 1E), whereas monensin treatment did not (Fig. 1F). This suggested an interesting possibility that there is a crosstalk from the mucin pathway to the TFE3 pathway.

### BG enhances transcription from the promoter of the human TFE3 gene

To reveal the molecular mechanism of how mucin-type Golgi stress activates the transcription of TFE3, we investigated the effect of BG treatment on transcription from the promoter of the human TFE3 gene. The [-2017 to +291] region of the human TFE3 gene (numbers indicate the position from the transcription start site) and its deletion constructs were ligated to the firefly luciferase gene and transfected into HT29 cells. HT29 cells were then treated with BG for 48 h, and the relative luciferase activity from these constructs was measured. We found that all deletion constructs shown in Fig. 2 had high activity of transcriptional induction upon BG treatment, suggesting that the enhancer element regulating transcriptional induction is contained within the [-12 to +19] region (Fig. 2D).

**Fig. 2.**
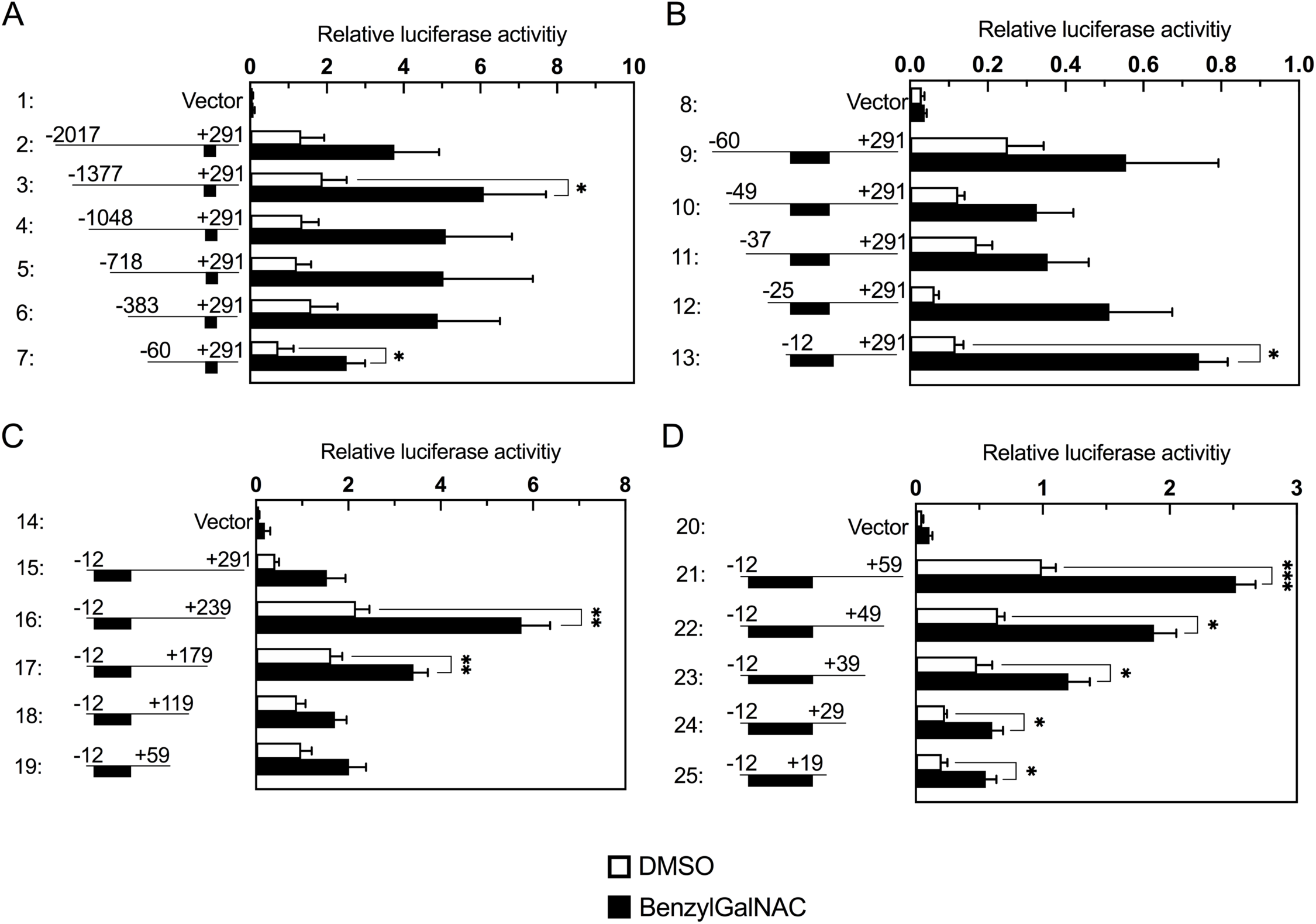
Promoter analysis of the human TFE3 promoter. HT29 cells were transiently transfected with reporter constructs containing the indicated regions of the human TFE3 promoter and a luciferase reporter gene, treated with 10 mM BG for 48 h, and subjected to luciferase assays. The estimated locations of enhancers regulating transcriptional induction are indicated by black boxes. The reporter plasmid pGL4 basic was used as control. Values are means ± SE of four independent experiments. ***, P < 0.001; **, P < 0.01; *, P < 0.05.

### Identification of an enhancer element activating transcriptional induction of the human TFE3 gene

To precisely delineate the nucleotide sequence of the enhancer element regulating the transcriptional induction of TFE3, we introduced point mutations for each nucleotide within the [-12 to +19] region and performed luciferase assays (Fig. 3). Compared with the luciferase activity of a wild-type (WT) sequence (lane 2), that of mutant sequences containing a single nucleotide mutation in Region A or Region B was considerably reduced (lanes 13-18 and lanes 28-33, respectively). These results suggested that the consensus sequence of the enhancer element is ACTTCC(N9)TCCCCA. We named this sequence the mucin-type Golgi stress response element (MGSE).

**Fig. 3.**
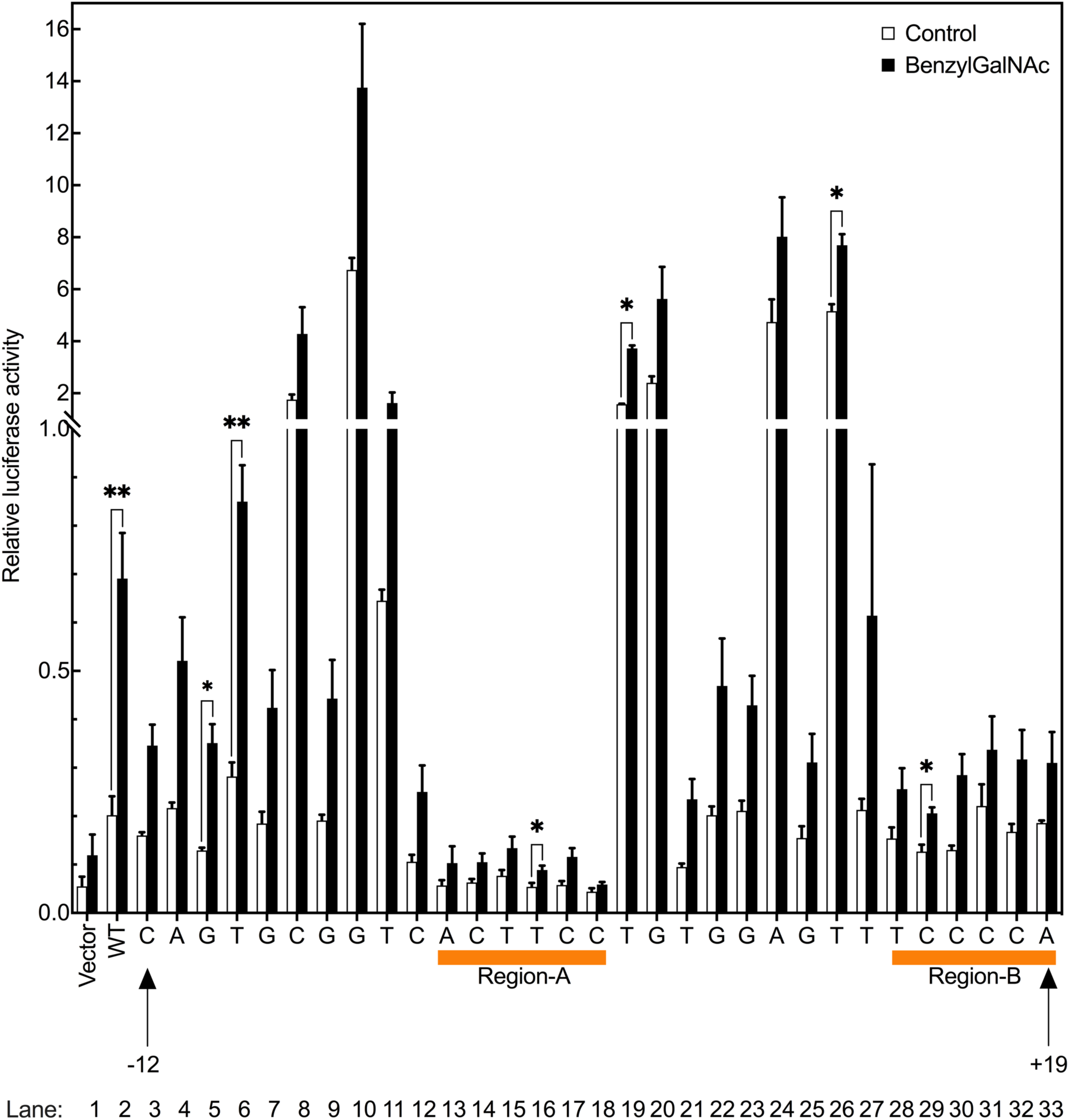
Point mutation analysis of the [-12 to +19] region of the human TFE3 promoter. HT29 cells were transiently transfected with a luciferase reporter containing the [-12 to +19] region of human TFE3 promoter, treated with 10 mM BG for 48 h, and subjected to luciferase assays. Each single nucleotide in the [-12 to +19] region was mutated to another nucleotide (A, T, G, and C were replaced by C, G, T, and A, respectively). The reporter plasmid pGL4 basic was used as vector control. Values are means ± SE of three independent experiments. ***, P < 0.001; **, P < 0.01; *, P < 0.05.

### The MGSE is necessary and sufficient for transcriptional induction of TFE3 upon BG treatment

To reveal whether the MGSE sequence is essential for transcriptional induction of the TFE3 promoter, we constructed mutant promoters of TFE3 in which the MGSE sequence was mutated (Fig. 4A). A vector control hardly responded to BG treatment (lane 1), whereas a reporter containing the WT promoter ([-12 to +292]) increased transcription of the luciferase gene after BG treatment (lane 2). When either the ACTTCC or TCCCCA motif was mutated, transcriptional induction was markedly reduced (lanes 3-5), indicating that both motifs of MGSE are essential for transcriptional induction of the TFE3 promoter.

**Fig. 4.**
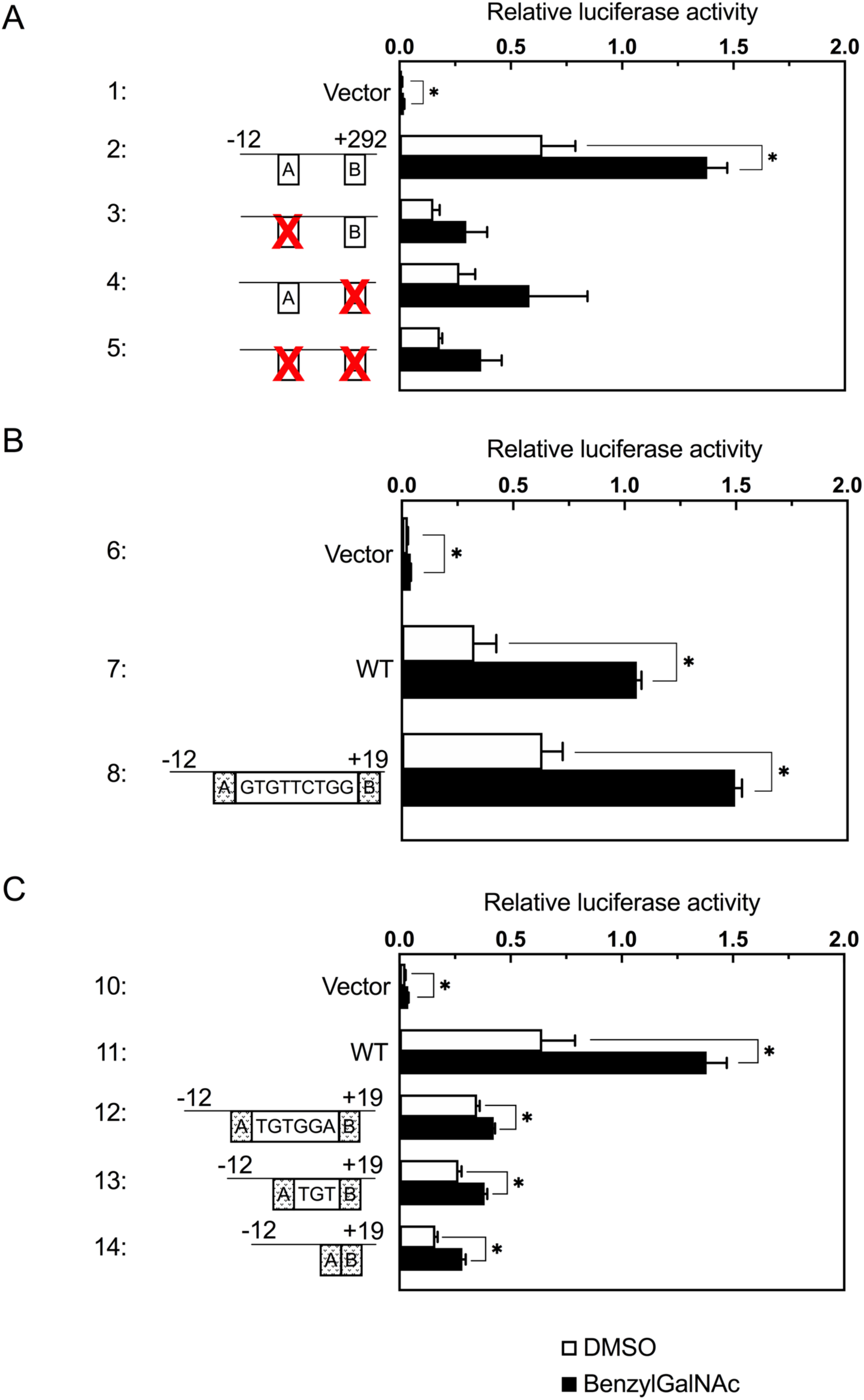
Contribution of MGSE to transcriptional induction of the TFE3 promoter and importance of the spacer region of MGSE. (A) Mutation of the MGSE sequence in the [-12 to +292] region of the human TFE3 promoter. HT29 cells transfected with the indicated reporter plasmids were treated with 10 mM BG for 48 h and subjected to luciferase assays. A and B indicate the ACTTCC and TCCCCA motifs of MGSE, respectively. (B) Mutation of the spacer sequence between ACTTCC and TCCCCA motifs of MGSE. The spacer sequence of wild-type ‘TGTGGAAGT’ was mutated to ‘GTGTTCTGG’. Activity of the mutated sequences ([-12 to +19]) were evaluated as described in (A). (C) Stepwise deletion mutation of the spacer sequence. Cells transfected with the indicated reporter plasmids were processed as described in (B). Reporter plasmid pGL4 basic was used as the control. Values are means ± SE of three independent experiments. ***, P < 0.001; **, P < 0.01; *, P < 0.05.

Next, we evaluated the importance of the spacer sequence “TGTGGAAGT” between the ACTTCC and TCCCCA motifs (Fig. 4B). A mutant MGSE, in which the spacer sequence was changed to “GTGTTCTGG” responded to BG treatment as well as that containing the WT spacer (lanes 7 and 8), suggesting that the nucleotide sequence of the spacer region itself is not essential for transcriptional induction.

Finally, we evaluated the importance of the 9 nt distance between the ACTTCC and TCCCCA motifs (Fig. 4C). Mutant MGSEs containing only 6, 3, or 0 nt spacer sequences showed considerably reduced activity (lanes 11-13), indicating that the 9 nt distance (but not the nucleotide sequence itself) is important. These results indicated that ACTTCC(N9)TCCCCA is the consensus sequence of MGSE.

### Transcription from MGSE is activated by overexpression of mucin core proteins

In the above experiments, we utilized BG to evoke mucin-type Golgi stress. However, it is possible that transcriptional induction from MGSE upon BG treatment is not mediated by mucin-type Golgi stress but by off-target effect of BG, because BG is a small chemical. To confirm that MGSE actually enhances transcription in response to mucin-type Golgi stress, we devised another method to evoke mucin-type Golgi stress, that is, overexpression of mucin core proteins such as MUC1 and MUC20. Mucins including MUC1 and MUC20 are modified with a number of mucin-type O-glycosylations in the Golgi apparatus (Jensen *et al.*, 2010). Thus, overexpression of MUC1 or MUC20 may cause insufficiency of glycosylation enzymes for mucins in the Golgi, resulting in the induction of mucin-type Golgi stress.

First, we confirmed that BG can disrupt Golgi morphology in HT29 cells. When HT29 cells were treated with BG and stained with anti-GM130 (a *cis*-Golgi marker) (Nakamura, 2010), the Golgi was highly distressed and disassembled (Fig. 5A, panels d-f), whereas the Golgi was intact in cells treated with DMSO, suggesting that BG induces Golgi stress and induces the disruption of Golgi morphology in HT29 cells. On the contrary, when HeLa cells, which synthesize lower amounts of mucin proteins, were treated with BG, disruption of the Golgi was less prominent (Fig. 5B). These results suggested that Golgi disruption induced by BG depends on mucin production. As for the reason why BG slightly disrupted the Golgi in HeLa cells with low mucin production, we speculated that HeLa cells synthesize mucin-type glycans, although we do not know if such mucin-type glycans are conjugated to mucin core proteins or other proteins. Proteins incompletely modified with mucin-type glycans might accumulate in the Golgi and cause disruption of Golgi morphology.

**Fig. 5.**
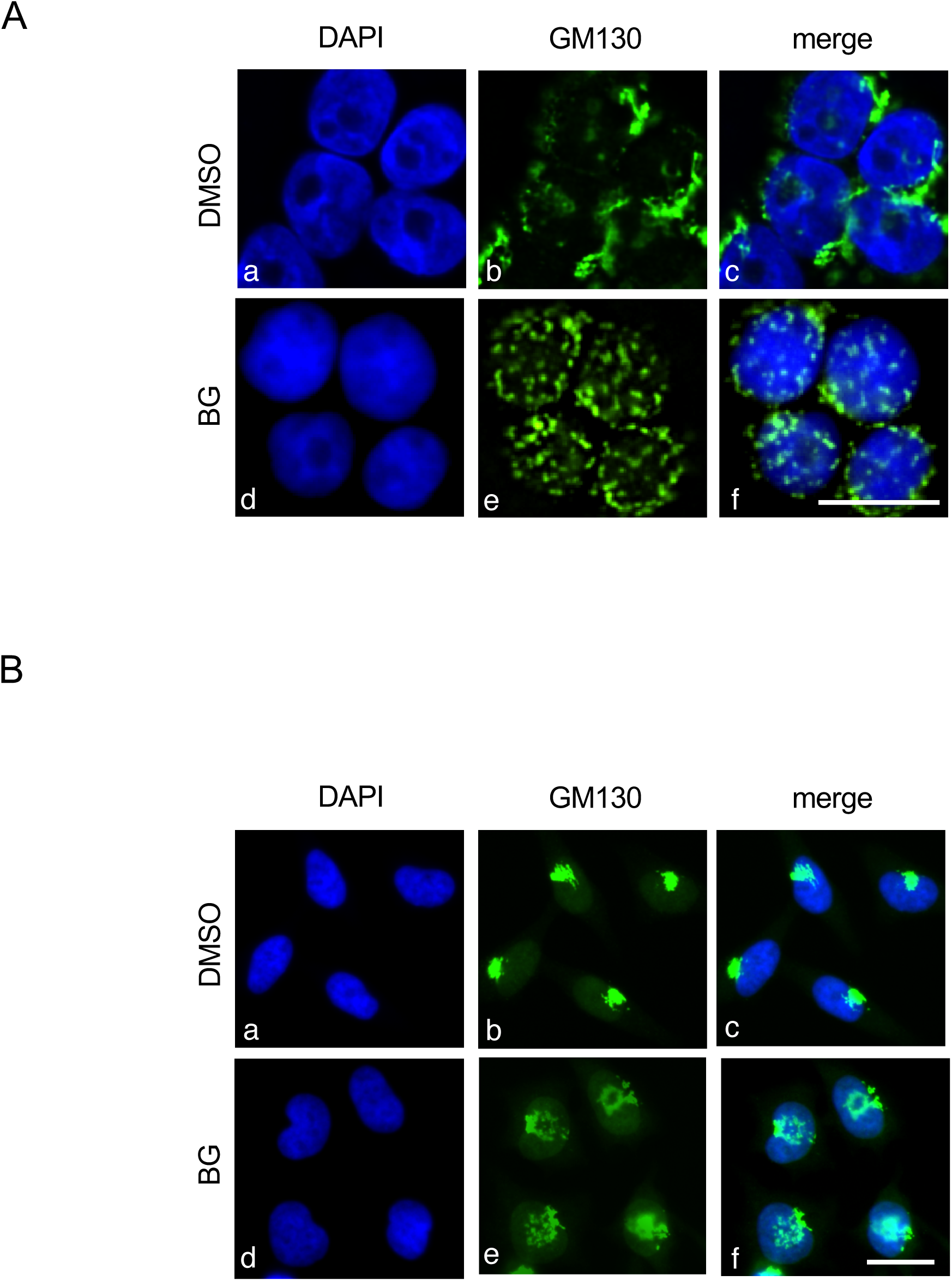
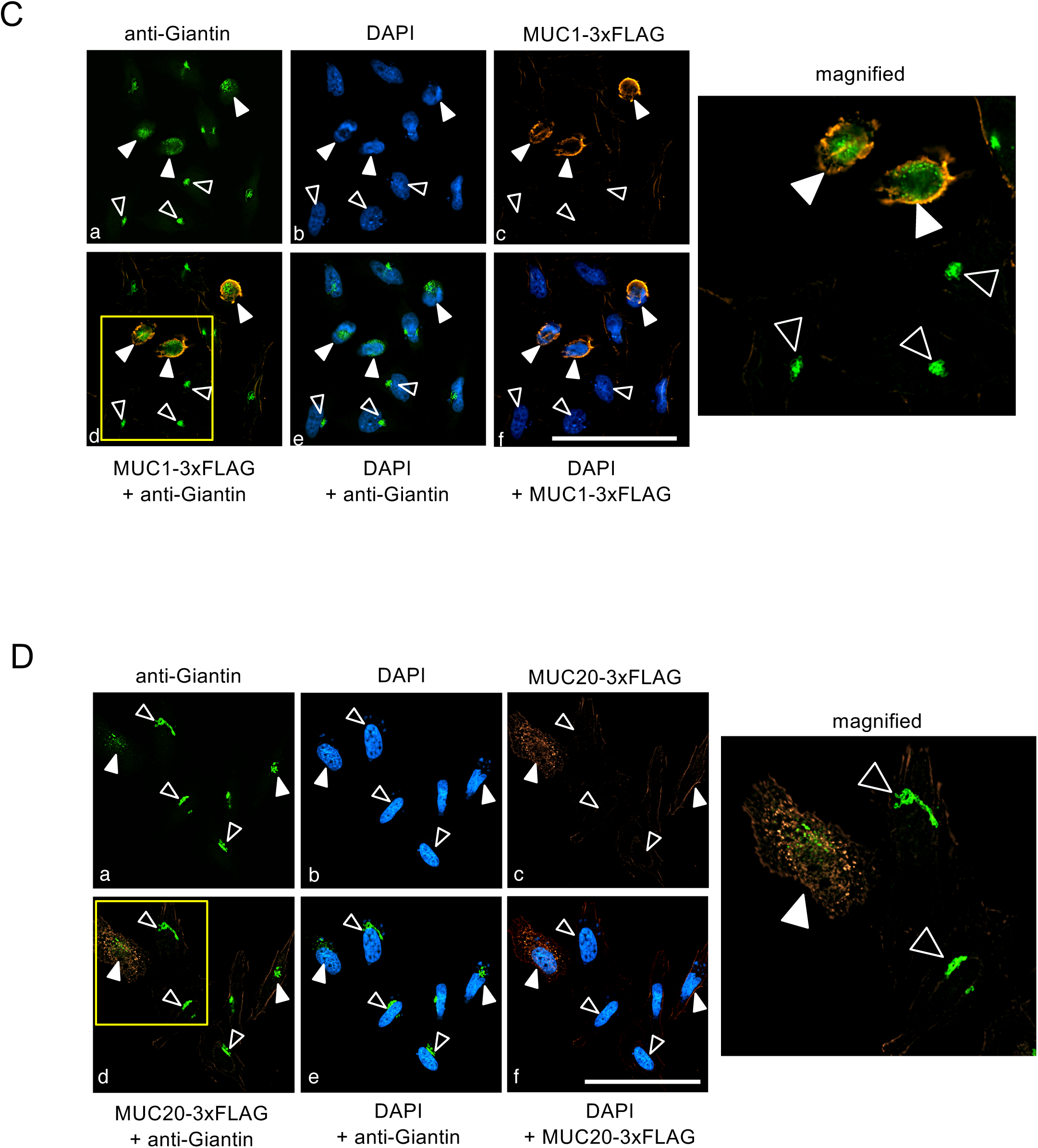
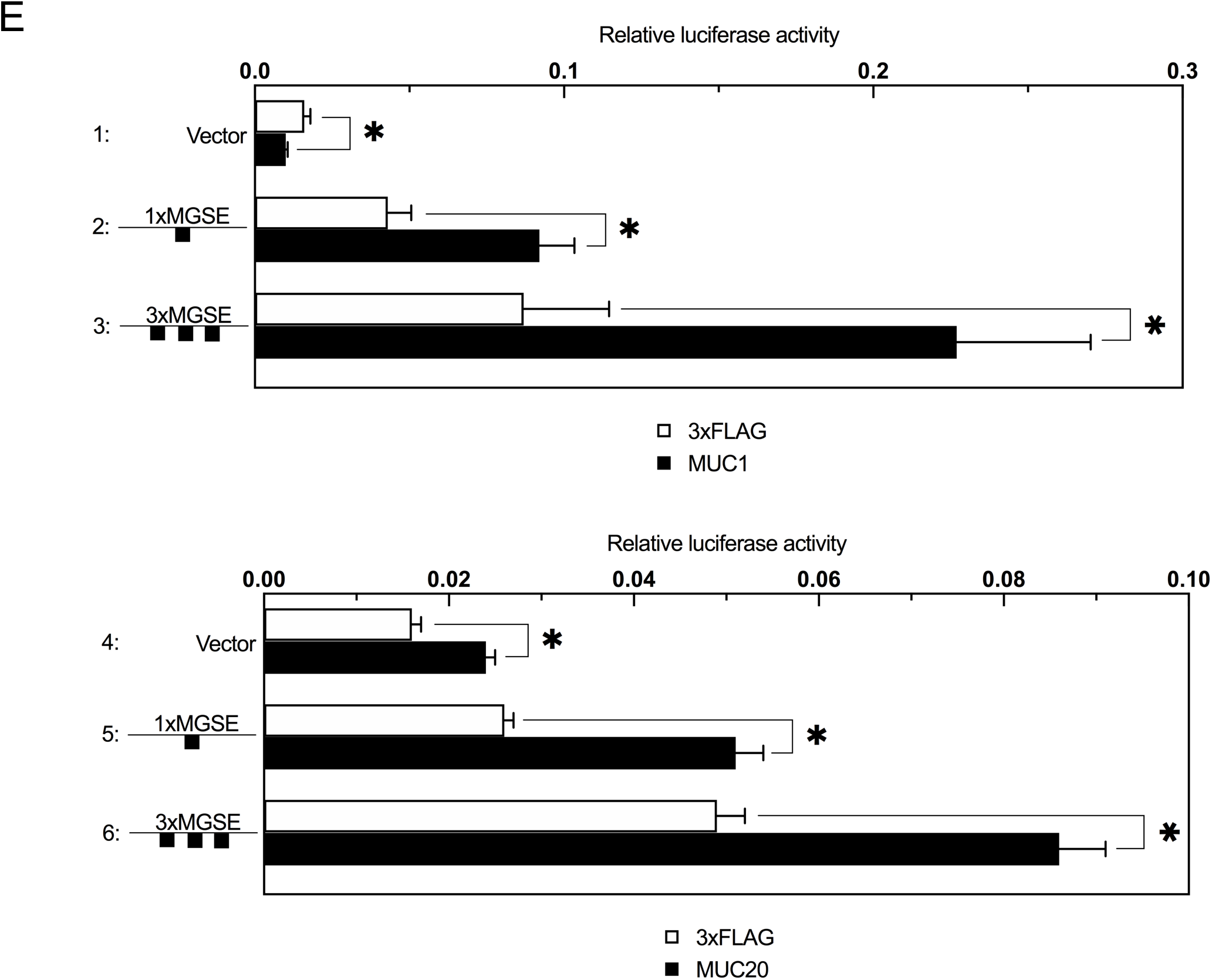
Morphological changes in the Golgi apparatus upon mucin type-Golgi stress. (A) HT29 cells treated with 10 mM BG for 48 h were stained with DAPI (blue) and anti-GM130 antibody (green). Bars = 10 μm. (B) The same experiment as (A) was performed using HeLa cells. (C) HeLa cells transfected with a 3×FLAG-MUC1 expression vector were stained with anti-FLAG (orange) and anti-Giantin (green) antisera. Bars = 50 μm. Closed and open arrow heads indicate cells that overexpressed or did not overexpressed 3×FLAG-MUC1, respectively. (D) HeLa cells transfected with a 3×FLAG-MUC20 expression vector were stained with anti-FLAG (orange) and anti-Giantin (green) antisera. Bars = 50 μm. Closed and open arrow heads indicate cells that overexpressed and did not overexpress 3×FLAG-MUC20, respectively. (E) HT29 cells transiently co-transfected with a 3×FLAG-MUC1 or 3×FLAG-MUC20 expression plasmid and a luciferase reporter containing 1×MGSE and 3×MGSE as indicated. Reporter plasmid pGL4 basic was used as a control. Values are means ± SE of three independent experiments. ***, P < 0.001; **, P < 0.01; *, P < 0.05.

Next, we examined the effect of MUC1 or MUC20 overexpression on the morphology of the Golgi (Fig. 5C and Fig 5D). For this experiments, we used HeLa cells instead of HT29 cells because HeLa cells synthesize lower amounts of mucins and the effect of overexpression of mucin core proteins in HeLa cells appeared larger than that in HT29 cells. HeLa cells were transfected with expression vectors 3×FLAG-MUC1 or 3×FLAG-MUC20 and stained with anti-Giantin (a *cis*- and *medial*-Golgi marker) (Sohda *et al.*, 2001) and anti-FLAG antisera. In HeLa cells that overexpressed 3×FLAG alone, the morphology of the Golgi was normal and intact (open arrow head), whereas the Golgi was fragmented and dispersed in cells overexpressing 3×FLAG-MUC1 or 3×FLAG-MUC20 (closed arrow heads), suggesting that overexpression of mucin core proteins overwhelmed the capacity of mucin-type glycosylation enzymes in the Golgi and induced mucin-type Golgi stress.

Finally, we evaluated the effect of MUC1 or MUC20 overexpression on transcription from MGSE (Fig. 5E). We co-transfected HT29 cells with an expression vector of 3×FLAG-MUC1 or 3×FLAG-MUC20 as well as a 1× or 3×MGSE reporter, and measured relative luciferase activity. In cells transfected with a control reporter without MGSE, overexpression of MUC1 or MUC20 hardly affected luciferase activity (lane 1 and lane 4 respectively), whereas luciferase activity was significantly increased by overexpression of MUC1 or MUC20 in cells transfected with MGSE reporters (lanes 2, 3, 5, and 6), suggesting that transcription from MGSE is activated by the overexpression of mucin core proteins. The above results strongly supported our notion that MGSE is an enhancer regulating transcriptional induction of TFE3 in response to mucin-type Golgi stress.

### Mucin-type Golgi stress evoked TFE3 dephosphorylation and nuclear translocation

Our finding that mucin-type Golgi stress induces the transcription of TFE3 suggested an interesting possibility that mucin-type Golgi stress may activate the TFE3 pathway. TFE3 is retained as a dormant form in the cytoplasm through phosphorylation at Ser108 in normal growth conditions, whereas upon activation of the TFE3 pathway by Golgi stress TFE3 is dephosphorylated, translocates into the nucleus, and activates transcription from GASE (Taniguchi et al., 2015). Thus, we sought to verify this notion by examining the activation status of the TFE3 pathway upon mucin-type Golgi stress.

First, we examined whether TFE3 translocates into the nucleus upon BG treatment (Fig. 6). When endogenous TFE3 was stained with anti-TFE3-A antiserum in control HT29 cells, TFE3 was observed in the cytoplasm (panels a–c). Upon BG treatment, TFE3 was localized in the nucleus (panels d–r), suggesting that TFE3 translocates upon mucin-type Golgi stress. We do not know why TFE3 in BG-treated cells was stained as particles in the nucleus, and we speculated that TFE3 was located on the promoters of its target genes or TFE3 formed liquid droplets by liquid–liquid phase separation. We then further examined TFE3 subcellular localization upon overexpression of MUC1 or MUC20 and found that TFE3 was localized in the nucleus in cells overexpressing MUC1 or MUC20 (closed arrow head) (Fig. 7A and 7B). On the other hand, TFE3 was localized in cytoplasm (open arrow head) in cells that did not overexpress 3×FLAG-MUC1 and 3×FLAG-MUC20 respectively, suggesting that mucin-type Golgi stress induces nuclear translocation of TFE3.

**Fig. 6.**
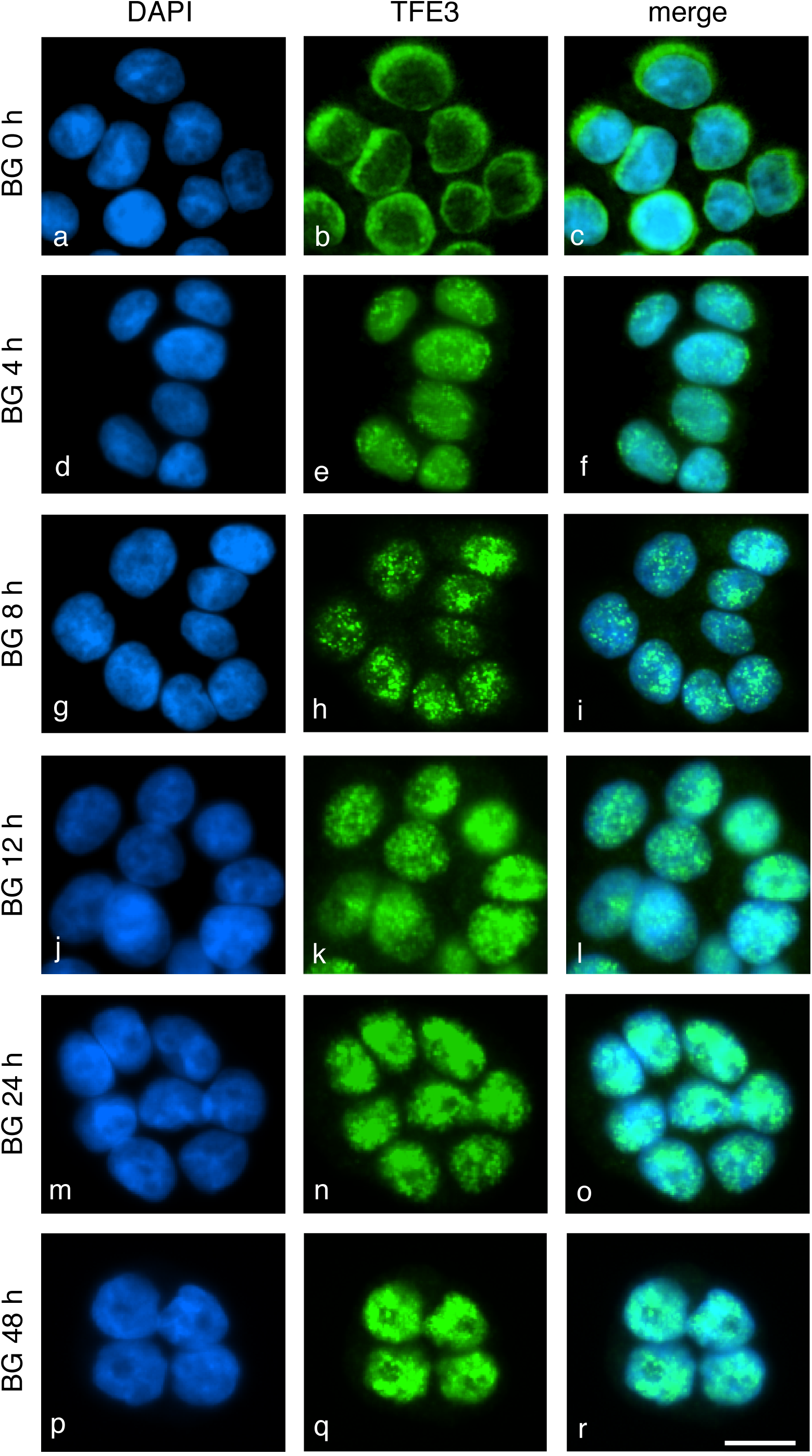
Nuclear translocation of TFE3 upon BG treatment. HT29 cells treated with 10 mM of BG for the indicated times were subjected to immunocytochemistry using anti-TFE3 antiserum and DAPI for staining. Bars = 20 μM.

**Fig. 7.**
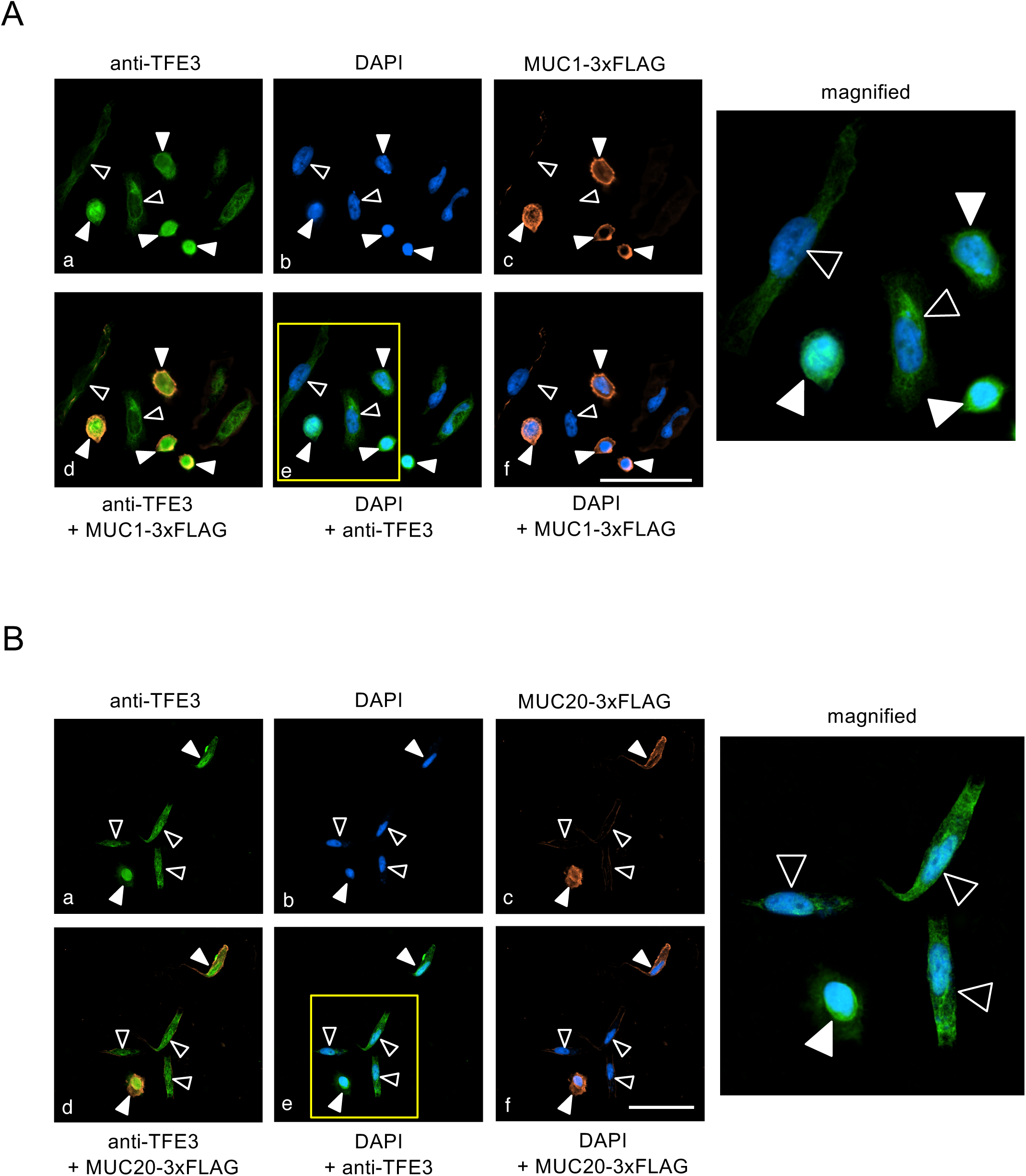
Nuclear translocation of TFE3 upon MUC1 or MUC20 overexpression. (A) HeLa cells transfected with a 3×FLAG-MUC1 expression vector were stained with anti-FLAG (orange) and anti-TFE3 (green) antisera. Bars = 50 μm. Closed and open arrow heads indicate cells that overexpressed or did not overexpress 3×FLAG-MUC1, respectively. (B) HeLa cells transfected with a 3×FLAG-MUC20 expression vector were stained with anti-FLAG (orange) and anti-TFE3 (green) antisera. Bars = 50 μm. Closed and open arrow heads indicate cells that overexpressed and did not overexpress 3×FLAG-MUC20, respectively.

Second, we investigated the phosphorylation status of TFE3 upon mucin-type Golgi stress (Fig. 8A and 8B). In the absence of Golgi stress, we observed eight bands of TFE3 as reported previously (lane 1) (Taniguchi et al., 2015). The four upper bands correspond to the large forms of TFE3 (TFE3(L)), and the four lower bands are derived from the short forms (TFE3(S)). The difference in molecular weight of each of the four bands is due to differences in phosphorylation status as shown in Fig. 8A. When HT29 cells were treated with vehicle (DMSO), the phosphorylation status of TFE3 hardly changed (lanes 2–4). In contrast, TFE3 was dephosphorylated 8 h after BG treatment (lanes 7 and 8), suggesting that mucin-type Golgi stress induces TFE3 dephosphorylation. Expression of TFE3 was increased 8 h after BG treatment, reflecting transcriptional induction of TFE3 by mucin-type Golgi stress.

**Fig. 8.**
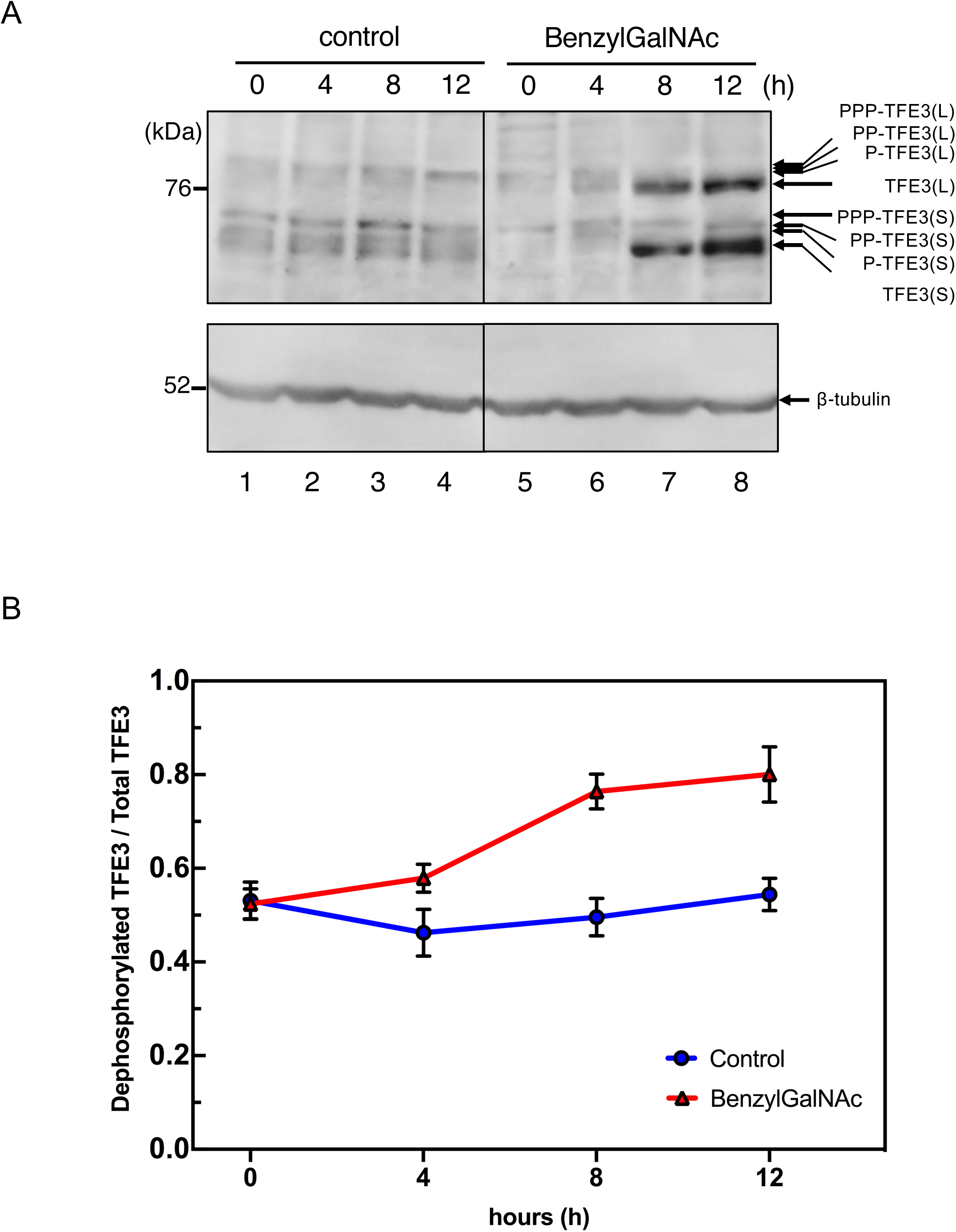
Phosphorylation status of TFE3 upon BG treatment. (A) HT29 cells treated with 10 mM BG for the indicated time periods were subjected to western blotting using anti-TFE3 and anti-β-tubulin antisera. TFE3(L) and (S) are long and short forms of TFE3, respectively, and (P-) represents phosphorylated TFE3. (B) Ratio of signal detected between dephosphorylated TFE3 and total TFE3. Signals of TFE3(L) and (S) in (B) were quantified and plotted accordingly.

Third, we evaluated transcriptional induction from GASE upon mucin-type Golgi stress (Fig. 9). As expected, overexpression of TFE3 increased transcription from GASE (lane 2, blue bar), but not from mutant GASE (lane 3, blue bar). Surprisingly, BG alone did not affect transcription from GASE. This meant that mucin-type Golgi stress increases transcription, nuclear translocation, and dephosphorylation of TFE3, which is not sufficient for transcriptional induction from GASE (see discussion).

**Fig. 9.**
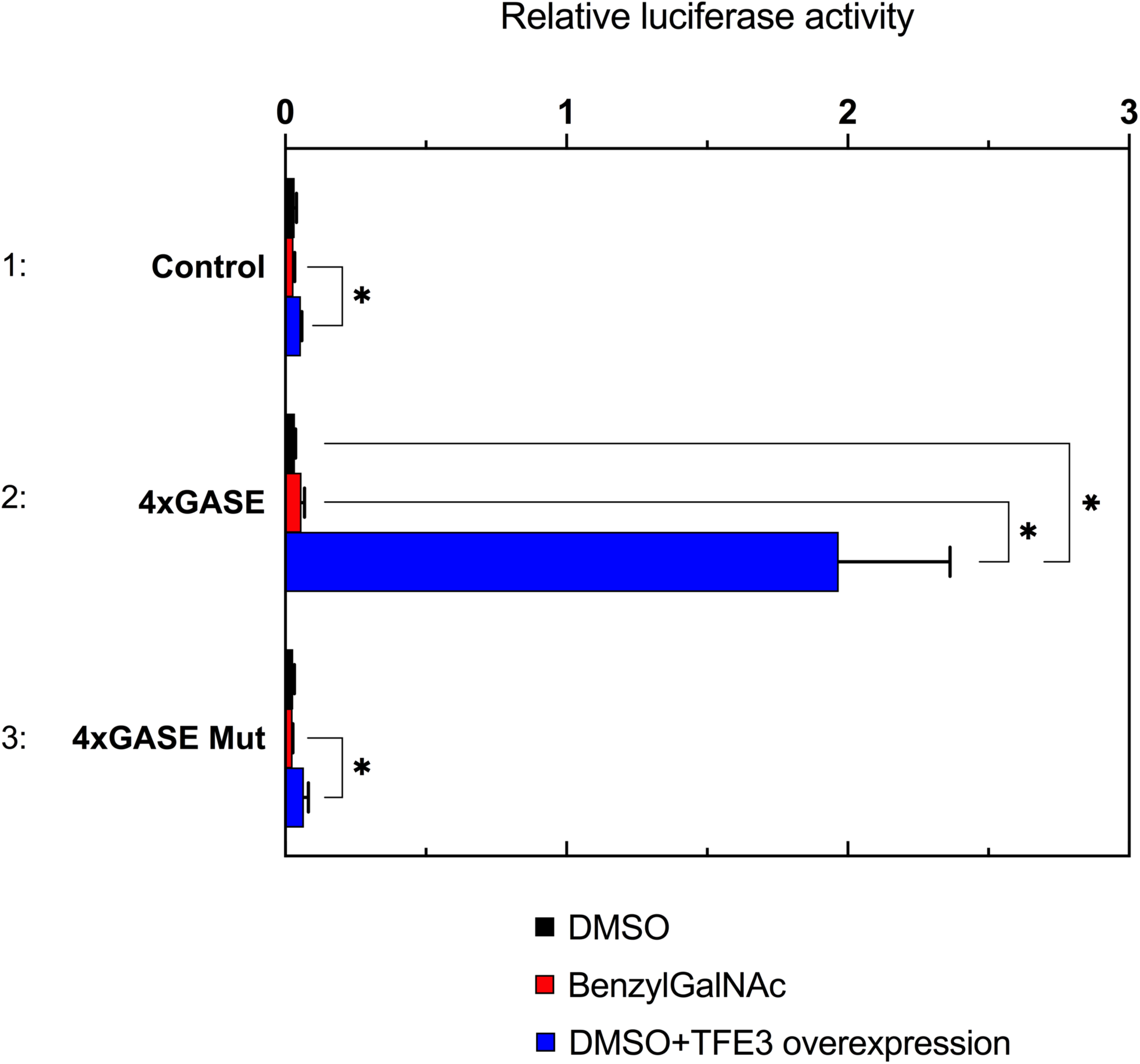
Effects of BG treatment on GASE-mediated transcriptional induction. HT29 cells were transiently transfected with the indicated plasmids and subjected to luciferase assays. Reporter plasmid pGL3pro was used as the control, and is shown in lane 1.

## Discussion

Mucin-type glycosylation of proteins is an important function of the Golgi apparatus, suggesting that mucin-secreting cells have to augment the expression of enzymes for mucin-type glycosylation via an unidentified pathway of the Golgi stress response. Here we found that the expression levels of GALNT5, GALNT8, and GALNT18 were increased upon insufficiency of mucin-type glycosylation (mucin-type Golgi stress), and we named this pathway the mucin pathway. Unexpectedly, we found that the expression of TFE3, a transcription factor regulating the TFE3 pathway of the Golgi stress response, was also increased in response to mucin-type Golgi stress, indicating the existence of crosstalk from the mucin pathway to the TFE3 pathway. Through analysis of the human TFE3 promoter and its mutants, we identified a novel enhancer regulating transcriptional induction of the TFE3 gene upon mucin-type Golgi stress, which we named MGSE, for which the consensus sequence was ACTTCC(N9)TCCCCA. We also found that TFE3 is dephosphorylated and translocates into the nucleus upon mucin-type Golgi stress, although transcription from GASE was not affected by mucin-type Golgi stress. From these observations, we concluded that the mucin pathway generates a crosstalk signal to the TFE3 pathway.

The distance between the ACTTCC and TCCCCA motifs was critical for the activity of MGSE. This may indicate that two distinct transcription factors synergistically bind to each portion. A similar situation is found in the case of ER stress response element (ERSE; consensus sequence CCAAT(N9)CCACG), an enhancer regulating the ATF6 pathway of the mammalian ER stress response (Yoshida *et al.*, 1998). The transcription factor NF-Y binds to the CCAAT sequence, and another transcription factor, ATF6, requires both the CCACG sequence and NF-Y binding to the CCAAT sequence for its binding. Because the distance of 9 nt almost corresponds to one turn of a DNA double helix, transcription factors binding to MGSE seem analogous the situation with ATF6 and NF-Y (Yoshida *et al.*, 2001).

We have been analyzing an enhancer element that regulates the transcriptional induction of GALNT5, GALNT8, and GALNT18 in response to mucin-type Golgi stress, and it is still unclear whether it is identical to MGSE. There did not appear to be MGSE-like sequences in the human GALNT5, GALNT8, and GALNT18 promoters, suggesting that the enhancer element is distinct from MGSE. As for an enhancer element regulating the transcriptional induction of HSP47 after BG treatment, we suspected that it is distinct from MGSE and the enhancer for GALNTs, because (1) there are no MGSE-like sequences in the human HSP47 promoter, and (2) the HSP47 pathway regulates a quite different response (not glycosylation but suppression of apoptosis) compared with the mucin pathway.

BG is distinct from authentic activators of the TFE3 pathway such as monensin and nigericin, because: (1) monensin does not affect the expression of TFE3 mRNA (Fig. 1F) (Taniguchi et al., 2015), whereas BG induces expression of the TFE3 mRNA (Fig. 1E); and (2) monensin activates the TFE3 pathway, including nuclear translocation and dephosphorylation of TFE3 as well as transcriptional induction from GASE (Oku et al., 2011), whereas BG induces nuclear translocation and dephosphorylation of TFE3 (Fig. 6) but does not increase transcriptional induction from GASE (Fig. 9). This indicated that BG does not activate a sensor for the TFE3 pathway, and that there are two kinds of crosstalk signals from the mucin pathway to the TFE3 pathway, namely a signal enhancing transcription of the TFE3 gene and a signal inducing dephosphorylation of TFE3 protein (Fig. 10). This raises the following three questions. First, why does BG alone not activate transcription from GASE? We speculated that it is because cells have to avoid accidental activation of the TFE3 pathway. If the mucin pathway directly activated transcription from GASE, it could be induced without activation of the TFE3 pathway. Second, how do cells suppress ectopic transcriptional activation from GASE upon BG treatment? It is notable that crosstalk signals induced by BG treatment increased expression and nuclear translocation of TFE3 proteins, but it was not enough for transcriptional activation from GASE. We speculated that the crosstalk signals may require additional signals in order to activate transcription from GASE, possibly additional post-translational modifications of TFE3 protein or transcription factors other than TFE3, which are induced or activated by the TFE3 pathway. Third, what is the biological significance of these crosstalk signals? We assumed that mucin-secreting cells require very high general Golgi capacity, such as vesicular transport and N-glycosylation, because they produce high numbers of mucin proteins. Thus, cells enhance the TFE3 pathway using these crosstalk signals during differentiation of mucin-secreting cells such as goblet cells.

**Fig. 10.**
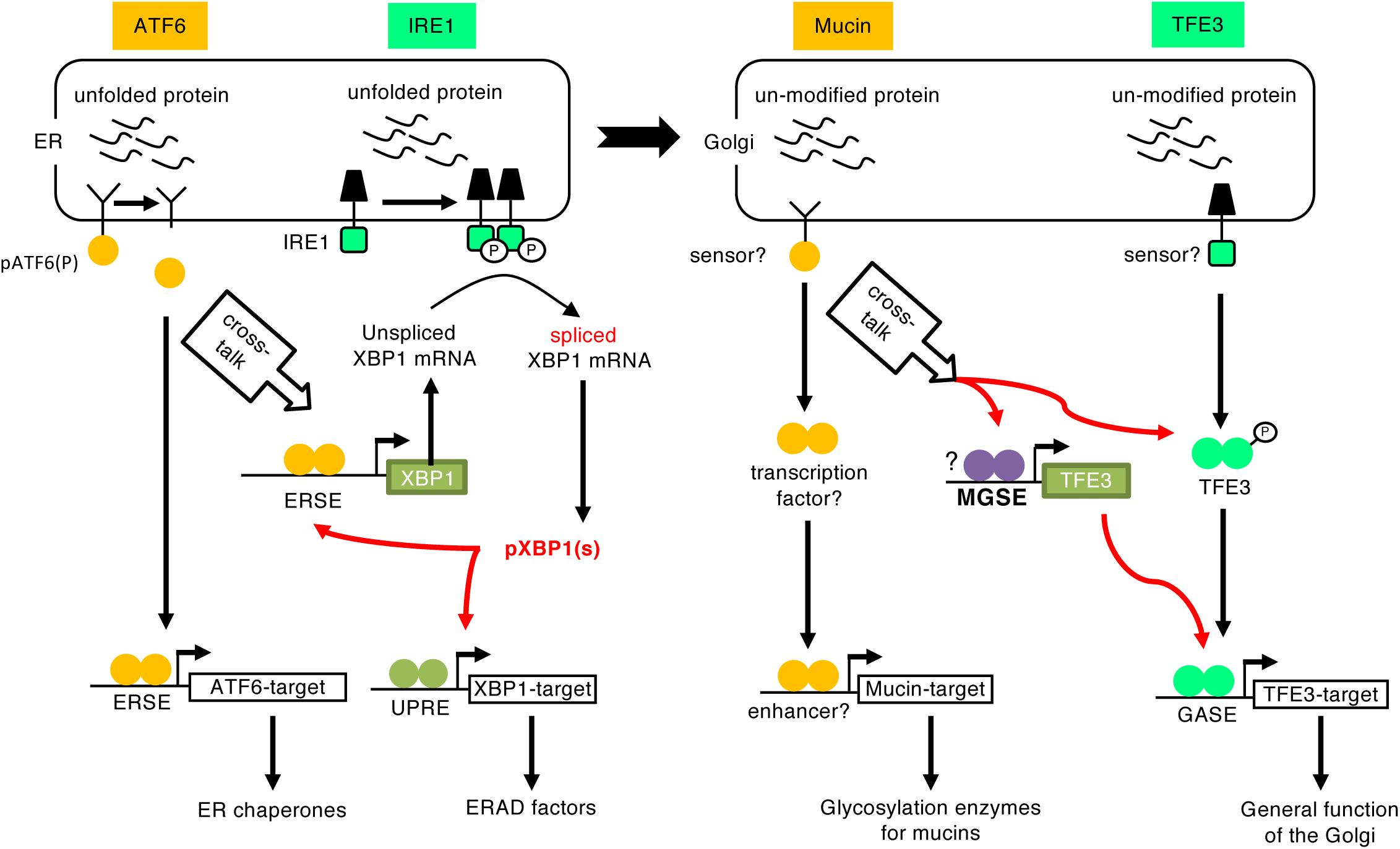
Working hypothesis of crosstalk signaling from the mucin pathway to the TFE3 pathway. Crosstalk from the ATF6 pathway to the IRE1 pathway in the ER stress response pathway is also indicated. A detailed explanation is given in the text.

In the case of ER stress response, crosstalk exists between the IRE1 and ATF6 pathways. pATF6(N), a transcription factor regulating the ATF6 pathway by upregulating transcription of ER chaperones (Yamamoto *et al.*, 2007), increases the transcription of XBP1 (Yoshida *et al.*, 2000), which is a key transcription factor upregulating the transcription of ERAD factors (Yoshida *et al.*, 2003). Thus, the ATF6 pathway enhances the response of the IRE1 pathway. However, the ATF6 pathway only partially activates the IRE1 pathway, because ATF6 just increases the expression of XBP1 pre-mRNA, which has to be converted to mature mRNA by IRE1-mediated non-conventional mRNA splicing in order to produce an active transcription factor pXBP1(S) (Fig. 10) (Yoshida et al., 2001).

Like the ER and Golgi stress responses, many studies are now progressing to clarify the mechanism that augments the capacity of each organelle in accordance with cellular demands, including the lysosomal stress response (Napolitano and Ballabio, 2016), mitochondrial unfolded protein response (Shpilka and Haynes, 2018), and peroxisome stress response, which are collectively called organelle autoregulation (Sasaki and Yoshida, 2015). Elucidation of the mechanisms of organelle autoregulation is one of the most fundamental issues in cell biology, because they are essential for the autonomy of eukaryotic cells. Insufficiency of functional capacity is sensed by sensors localized on or in each organelle, which relay signals to transcription factors to augment their functions. Subsequently, transcription factors bind to unique enhancer elements to increase the expression of organelle-specific genes. In order to understand the perspective of organelle autoregulation, clarification of these regulators is crucial.

Here we identified MGSE and highlighted the existence of crosstalk among mammalian Golgi stress response pathways, which will contribute to identifying the sensor and the transcription factor. Our findings make a promising foundation in revealing the whole the Golgi stress response mechanism and support medical research on Golgi-associated diseases.

## Acknowledgement

We thank Ms. Mikiko Ochiai for secretarial assistance. This work was supported by JSPS KAKENHI (Grant numbers JP16K07356, JP15J05492, JP17K15122 and JP17J00067, a Grant-in-Aid for Scientific Research on Innovative Areas of MEXT (JP17H06414) and JSPS Bilateral Joint Research Projects.

## References

Gething, M.-J. 1997. Guidebook to Molecular Chaperones and Protein-Folding Catalysts. Oxford University Press, Oxford. pp.

Huet, G., Hennebicq-Reig, S., de Bolos, C., Ulloa, F., Lesuffleur, T., Barbat, A., Carriere, V., Kim, I., Real, F.X., Delannoy, P., and Zweibaum, A. 1998. GalNAc-alpha-O-benzyl inhibits NeuAcalpha2-3 glycosylation and blocks the intracellular transport of apical glycoproteins and mucus in differentiated HT-29 cells. J. Cell Biol., 141: 1311–1322.

Jensen, P.H., Kolarich, D., and Packer, N.H. 2010. Mucin-type O-glycosylation--putting the pieces together. FEBS J, 277: 81–94.

Karagoz, G.E., Acosta-Alvear, D., and Walter, P. 2019. The Unfolded Protein Response: Detecting and Responding to Fluctuations in the Protein-Folding Capacity of the Endoplasmic Reticulum. Cold Spring Harb. Perspect. Biol.

Kimata, Y. and Kohno, K. 2011. Endoplasmic reticulum stress-sensing mechanisms in yeast and mammalian cells. Curr. Opin. Cell Biol., 23: 135–142.

Komori, R., Taniguchi, M., Ichikawa, Y., Uemura, A., Oku, M., Wakabayashi, S., Higuchi, K., and Yoshida, H. 2012. Ultraviolet a induces endoplasmic reticulum stress response in human dermal fibroblasts. Cell Struct. Funct., 37: 49–53.

Kuan, S.F., Byrd, J.C., Basbaum, C., and Kim, Y.S. 1989. Inhibition of mucin glycosylation by aryl-N-acetyl-alpha-galactosaminides in human colon cancer cells. The Journal of biological chemistry, 264: 19271–11927.

Miyata, S., Mizuno, T., Koyama, Y., Katayama, T., and Tohyama, M. 2013. The endoplasmic reticulum-resident chaperone heat shock protein 47 protects the Golgi apparatus from the effects of O-glycosylation inhibition. PLoS One, 8: e69732.

Mori, K. 2015. The unfolded protein response: the dawn of a new field. Proc. Jpn. Acad. Ser. B Phys. Biol. Sci., 91: 469–480.

Nakamura, N. 2010. Emerging new roles of GM130, a cis-Golgi matrix protein, in higher order cell functions. J. Pharmacol. Sci., 112: 255–264.

Napolitano, G. and Ballabio, A. 2016. TFEB at a glance. J. Cell Sci., 129: 2475–2481.

Oku, M., Tanakura, S., Uemura, A., Sohda, M., Misumi, Y., Taniguchi, M., Wakabayashi, S., and Yoshida, H. 2011. Novel cis-acting element GASE regulates transcriptional induction by the Golgi stress response. Cell Struct. Funct., 36: 1–12.

Reiling, J.H., Olive, A.J., Sanyal, S., Carette, J.E., Brummelkamp, T.R., Ploegh, H.L., Starnbach, M.N., and Sabatini, D.M. 2013. A CREB3-ARF4 signalling pathway mediates the response to Golgi stress and susceptibility to pathogens. Nat. Cell Biol., 15: 1473–1485.

Sakamoto, K. and Kadomatsu, K. 2017. Mechanisms of axon regeneration: The significance of proteoglycans. Biochim. Biophys. Acta, 1861: 2435–2441.

Sasaki, K., Komori, R., Taniguchi, M., Shimaoka, A., Midori, S., Yamamoto, M., Okuda, C., Tanaka, R., Sakamoto, M., Wakabayashi, S., and Yoshida, H. 2019. PGSE Is a Novel Enhancer Regulating the Proteoglycan Pathway of the Mammalian Golgi Stress Response. Cell Struct. Funct., 44: 1–19.

Sasaki, K. and Yoshida, H. 2015. Organelle autoregulation-stress responses in the ER, Golgi, mitochondria and lysosome. J. Biochem., 157: 185–195.

Sasaki, K. and Yoshida, H. 2019. Organelle Zones. Cell Struct. Funct.: in press.

Shpilka, T. and Haynes, C.M. 2018. The mitochondrial UPR: mechanisms, physiological functions and implications in ageing. Nat. Rev. Mol. Cell Biol., 19: 109–120.

Sohda, M., Misumi, Y., Yamamoto, A., Yano, A., Nakamura, N., and Ikehara, Y. 2001. Identification and characterization of a novel Golgi protein, GCP60, that interacts with the integral membrane protein giantin. J. Biol. Chem., 276: 45298–45306.

Taniguchi, M., Nadanaka, S., Tanakura, S., Sawaguchi, S., Midori, S., Kawai, Y., Yamaguchi, S., Shimada, Y., Nakamura, Y., Matsumura, Y., Fujita, N., Araki, N., Yamamoto, M., Oku, M., Wakabayashi, S., Kitagawa, H., and Yoshida, H. 2015. TFE3 Is a bHLH-ZIP-type Transcription Factor that Regulates the Mammalian Golgi Stress Response. Cell Struct. Funct., 40: 13–30.

Taniguchi, M., Sasaki-Osugi, K., Oku, M., Sawaguchi, S., Tanakura, S., Kawai, Y., Wakabayashi, S., and Yoshida, H. 2016. MLX Is a Transcriptional Repressor of the Mammalian Golgi Stress Response. Cell Struct. Funct., 41: 93–104.

Taniguchi, M. and Yoshida, H. 2017. TFE3, HSP47, and CREB3 Pathways of the Mammalian Golgi Stress Response. Cell Struct. Funct., 42: 27–36.

Uemura, A., Taniguchi, M., Matsuo, Y., Oku, M., Wakabayashi, S., and Yoshida, H. 2013. UBC9 Regulates the Stability of XBP1, a Key Transcription Factor Controlling the ER Stress Response. Cell Struct. Funct., 38: 67–79.

Volmer, R. and Ron, D. 2015. Lipid-dependent regulation of the unfolded protein response. Curr. Opin. Cell Biol., 33: 67–73.

Wu, X. and Rapoport, T.A. 2018. Mechanistic insights into ER-associated protein degradation. Curr Opin Cell Biol., 53: 22–28.

Yamamoto, K., Sato, T., Matsui, T., Sato, M., Okada, T., Yoshida, H., Harada, A., and Mori, K. 2007. Transcriptional induction of mammalian ER quality control proteins is mediated by single or combined action of ATF6alpha and XBP1. Dev. Cell, 13: 365–376.

Yoshida, H. 2007. ER stress and diseases. FEBS J, 274: 630–658.

Yoshida, H., Haze, K., Yanagi, H., Yura, T., and Mori, K. 1998. Identification of the cis-acting endoplasmic reticulum stress response element responsible for transcriptional induction of mammalian glucose-regulated proteins. Involvement of basic leucine zipper transcription factors. J. Biol. Chem., 273: 33741–33749.

Yoshida, H., Matsui, T., Hosokawa, N., Kaufman, R.J., Nagata, K., and Mori, K. 2003. A time-dependent phase shift in the mammalian unfolded protein response. Dev. Cell, 4: 265–271.

Yoshida, H., Matsui, T., Yamamoto, A., Okada, T., and Mori, K. 2001. XBP1 mRNA is induced by ATF6 and spliced by IRE1 in response to ER stress to produce a highly active transcription factor. Cell, 107: 881–891.

Yoshida, H., Okada, T., Haze, K., Yanagi, H., Yura, T., Negishi, M., and Mori, K. 2000. ATF6 activated by proteolysis binds in the presence of NF-Y (CBF) directly to the cis-acting element responsible for the mammalian unfolded protein response. Mol. Cell. Biol., 20: 6755–6767.

Yoshida, H., Okada, T., Haze, K., Yanagi, H., Yura, T., Negishi, M., and Mori, K. 2001. Endoplasmic reticulum stress-induced formation of transcription factor complex ERSF including NF-Y (CBF) and activating transcription factors 6alpha and 6beta that activates the mammalian unfolded protein response. Mol. Cell. Biol., 21: 1239–1248.

Yoshida, H., Oku, M., Suzuki, M., and Mori, K. 2006. pXBP1(U) encoded in XBP1 pre-mRNA negatively regulates unfolded protein response activator pXBP1(S) in mammalian ER stress response. J. Cell Biol., 172: 565–575.

Yoshida, H., Uemura, A., and Mori, K. 2009. pXBP1(U), a negative regulator of the unfolded protein response activator pXBP1(S), targets ATF6 but not ATF4 in proteasome-mediated degradation. Cell Struct. Funct., 34: 1–10.

